# A METHOD FOR INTEGRATIVE ANALYSIS OF LOCAL AND GLOBAL BRAIN DYNAMICS

**DOI:** 10.1101/2021.05.18.444698

**Authors:** Robyn L. Miller, Victor M. Vergara, Vince D. Calhoun

## Abstract

The most common pipelines for studying time-varying network connectivity in resting state functional magnetic resonance imaging (rs-fMRI) operate at the whole brain level, capturing a small discrete set of “states” that best represent time-resolved joint measures of connectivity over all network pairs in the brain. This whole-brain hidden Markov model (HMM) approach “uniformizes” the dynamics over what is typically more than 1000 pairs of networks, forcing each time-resolved high-dimensional observation into its best-matched high-dimensional state. While straightforward and convenient, this HMM simplification obscures functional and temporal nonstationarities that could reveal systematic, informative features of resting state brain dynamics at a more granular scale. We introduce a framework for studying functionally localized dynamics that intrinsically embeds them within a whole-brain HMM frame of reference. The approach is validated in a large rs-fMRI schizophrenia study where it identifies group differences in localized patterns of entropy and dynamics that help explain consistently observed differences between schizophrenia patients and controls in occupancy of whole-brain dFNC states more mechanistically.

## 1. INTRODUCTION

Brain network connectivity in resting-state functional magnetic resonance imaging (rs-fMRI) was, until recently, studied primarily as a stationary property assessed over the full duration of the scan. Working with time-varying measures of resting state connectivity, so-called *dynamic functional network connectivity* (dFNC) has gained traction in recent years, with much of this work based on a pipeline introduced in [1]. The key features of this approach are that it uses functional networks spanning the whole brain, estimates time-varying connectivity between the networks using correlations computed on sliding windows through the network timeseries and then clusters the resulting collection of high-dimensional observations into states (cluster centroids) that represent replicable, transiently realized whole-brain connectivity patterns. The number of unique networks pairs typically exceeds one thousand, while the optimal number of clusters according to standard criteria (e.g., elbow, silhouette) is typically less than eight. Standard measures of interest are the occupancy rate (OCR) and mean dwell time (MDT) of the cluster states. Transition probabilities between cluster states are also studied [2]. Given the prevalence of this general approach, addressing some of its limitations within its own terms, i.e., in a methodological ecosystem still framed by whole-brain HMM allows deeper information to emerge in a way that remains commensurable with earlier published studies. While the HMM simplification has several important limitations as a model for rs-fMRI, our focus here is on better resolving functionally localized features of the dynamic whole-brain connectome. We introduce a method that captures more granular, mechanistic processes of integrative and dissipative functional cohesion that undoubtedly drive much of the high-level dynamics observed with whole-brain HMM analysis. Moreover, we find that at this more granular level, the properties characteristic of schizophrenia patients also characterizes timepoints preceding transitions into whole-brain HMM states more occupied by patients, suggesting a lower-level mechanistic explanation for elevated SZ occupancies of these target HMM states.

## 2. METHODS

### 2.1 Data

We use data from a large, eyes-open resting-state fMRI study with approximately equal numbers of schizophrenia patients (SZs) and healthy controls (HCs) (*n* =311, nSZ=150). Imaging data for six of the seven sites was collected on a 3T Siemens Tim Trio System and on a 3T General Electric Discovery MR750 scanner at one site. The data was preprocessed with a standard, already published [2], pipeline and decomposed with group independent component analysis (GICA) into 100 group-level functional network spatial maps (SMs) and corresponding subject-specific timecourses (TCs). Through a combination of automated and manual pruning, *N*=47 functionally identifiable networks are retained. Subject specific spatial maps and timecourses were obtained from the group level spatial maps via spatio-temporal regression. The timecourses were detrended, despiked and subjected to some additional postprocessing steps. The networks obtained fall into 7 functional domains: subcortical (*SC, 5 nodes*), auditory (*AUD, 2 nodes*), visual (*VIS, 11 nodes*), sensorimotor (*SM, 6 nodes*), cognitive control (*CC, 13 nodes*), default mode (*DMN, 8 nodes*) and cerebellar (*CB, 2 nodes*). Functional domains are displayed along axes of connectivity matrices in the order given above. All subjects signed informed consent forms.

### 2.2 Dynamic Functional Network Connectivity

Dynamic functional connectivity (*dFNC*) between RSN timecourses was estimated using sliding window correlations. Following protocols from recent studies [2], dynamic functional network connectivity (*dFNC*) was estimated using pairwise correlations between RSN timecourses on tapered sliding rectangular windows of length 22 TRs (44 seconds), advancing TR at each step [2]. After dropping the first 3 TRs, this procedure yields a 44(47 − 1)/2=1081-dimensional dFNC measure on each of 136 windows of length 22 TRs for each subject. Using Matlab’s implementation of k-means, the resulting observations are placed into 5 clusters (elbow criterion, squared Euclidean metric, 2000 iterations, 500 replicates).

### 2.3 Elementwise Discrete Recoding of dFNCs

In addition to the cluster membership of each 1081-dimensional observation determined by k-means, we identify each element (or *cell*) of the dFNC observation with one of the five whole-brain clusters based on its *L*^2^ distance from the corresponding cell of each cluster centroid, 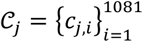, *j=* 1,2,..,5. Specifically, we apply a transformation *f*:[−1,1] → 𝒜 from the closed unit interval into the alphabet 𝒜=Error! Bookmark not defined. to each element *v*_*i*_ (*t*) ∈ [−1,1] of the observed dFNC matrix at time *t*, such that *f*(*v*_*i*_ (*t*)) = *α*_*i*_(*t*) ∈ 𝒜, where 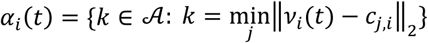 is the index of the whole-brain cluster whose corresponding element is closest to *v*_*i*_ (*t*). (**Figure 1**). whole-brain observation can then be considered either as belonging to its originally assigned cluster, or to the cluster to which the largest percentage of its elements are assigned (**Figure 1**). More importantly, this transformation allows domain blocks within the whole-brain connectome to be assigned a cluster in the same way, e.g., the CC-DMN block at time belongs to the cluster that the largest percentage of its elements are assigned to (**Figure 1**). The cellwise re-coded, 𝒜-valued, dFNCs are referred to as CCdFNCs.

**Figure 1.**
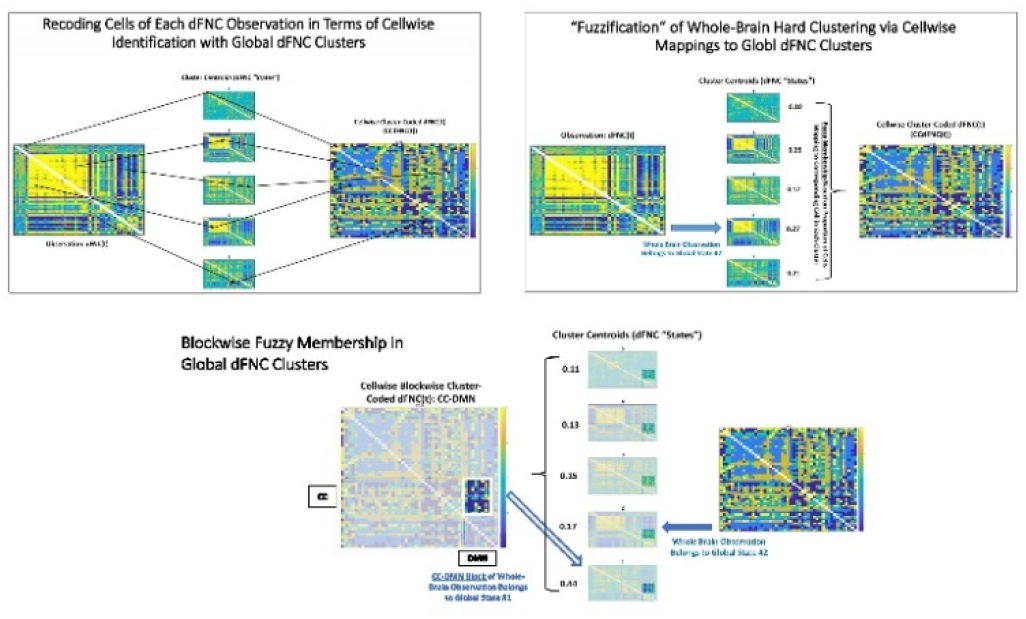
Schematic for cellwise recoding of dFNCs; (Top Left) Each cell of the observation is coded with the index of the whole-brain cluster to which the corresponding cell in the cluster centroid the observed value is closest to; (Top Right) This induces a fuzzy clustering of the dFNC based on the percentage of cells assigned to each whole-brain cluster; (Bottom) Domain blocks within the recoded dFNC are assigned cluster membership according to the percentage of cells in that block which are closest to the corresponding cell in each whole-brain cluster.

### 2.4 Time-Resolved and Dynamic Blockwise Entropy

Consisting of values from a finite alphabet, CCdFNCs offer a straightforward way to examine functional entropy across the whole connectome and localized to specific domain blocks (**Figure 2**). As a metric of the relative strength of integrative functional cohesion versus dissipative disorder, we compute the empirically observed entropy of the whole brain at time *t* as 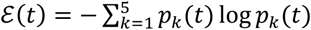 where *p*_*k*_= |{*α*_*i*_ (*t*) ∈ CCdFNC(*t*): *α*_*i*_ (*t*) =*k*}| / |CCdFNC |. Similarly, on the domain block level, the functionally localized time-resolved entropy (*tr*Ent) in block *B* ∈ CCdFNC at time *t* is 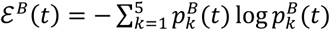 where 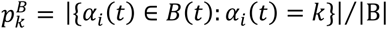. In the two degenerate blocks (AUD-AUD and CB-CB) with just one cell, ε^*B*^(*t*) ≡= 0. The AUD-CB block contains only 4 elements and thus cannot achieve the maximum possible entropy (− log(0.2) ≈ 1.61) on the five-letter alphabet 𝒜. They are included only for completeness and to maintain the structural shape of the connectome.

**Figure 2.**
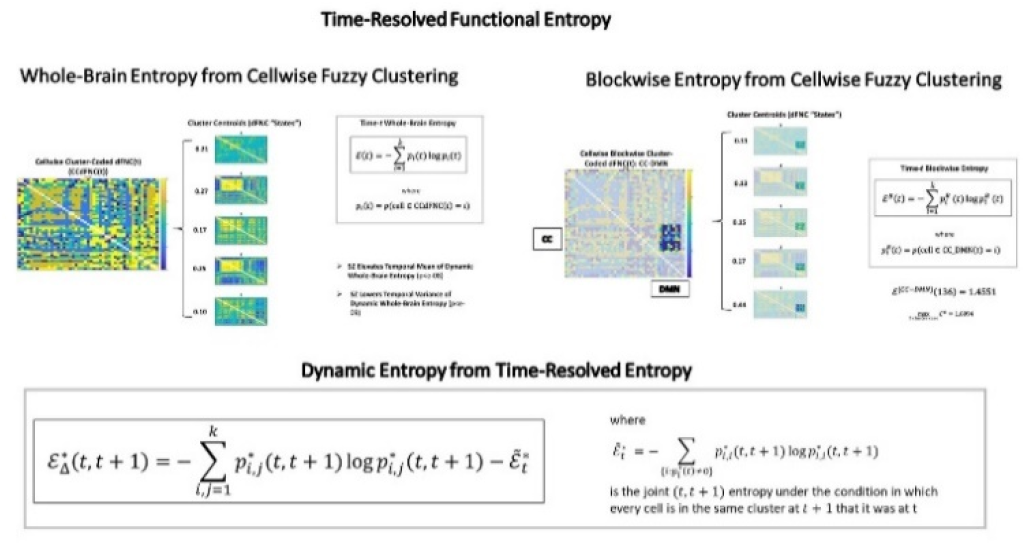
Whole brain (top left) and blockwise (top right) functional entropy is computed with respect to the observed probabilities of each value from the alphabet 𝒜 = {1,2,3,4,5} occurring in whole CCdFNC or in the block under consideration. Dynamic entropy (bottom) is computed with respect to observed probabilities of ordered pairs (*i,j*) from 𝒜 × 𝒜 occurring at consecutive timepoints *t, t* + 1 *corrected for* the entropy that would be observed if no cell changed assignment between *t* and *t* + 1.

To better resolve how integration and disorder evolve on short timescales, we also consider the *t* to *t* + 1 *dynamic* entropy (*dyn*Ent) on whole-brain and blockwise levels. In this case, the set of outcomes are the set of ordered pairs (*k,k*′) ∈ 𝒜 × 𝒜. The hypothetical case where *k* ′ = *k*, for all cells is the outcome in which none of the measured entropy is dynamic. Thus, we estimate the dynamic entropy on *B*(*t*) or CCdFNC(*t*) as 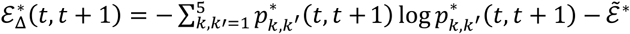 where 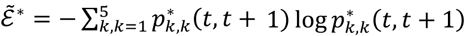.

### 2.5 Statistical Modeling

Reported SZ effects are from a multiple regression on gender, age, head motion and diagnosis. The effect of a condition applicable only to a subset of observations, e.g. observations at times {*t*:dFNC(*t*) is in whole brain cluster *k*}, on some measure *m* are estimated with *t*-tests of the measure taken under the condition of interest with respect to 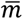, the global mean of the same measure. Results are only reported when significant at the *p* < 0.05 level after correction for multiple comparisons.

## 3. RESULTS

Using the discrete cellwise recoding of dFNC observations, we obtain insights into both global and functionally localized patterns of functional integrative and dissipative tendencies, and connections between these tendencies and diagnosed schizophrenia.

One important finding is that, not surprisingly, the auditory-visual-sensorimotor (AVSM) block – a very high magnitude, positively connected set of networks – drives whole-brain clustering to a significant degree. In particular, the visual block (**Figure 4**, left) is much more likely to belong to the cluster that k-means initially assigned the ambient whole-brain dFNC to. Other blocks have odds not far from chance (0.2) of being assigned to the “correct” cluster. This suggests that more sophisticated clustering strategies might be warranted for whole-brain analysis. The directional tendencies toward correct domain block assignment are amplified in the patient population (**Figure 4**, right). Which is to say that the whole-brain clusters are even less accurate descriptions of non-AVSM domains for SZs than for HCs.

We also find both whole-brain time-resolved entropy (*β*_*sz*_ =0.013, *p*_*sz*_ =2.5e^-08^) and whole-brain dynamic entropy (*β*_*sz*_ = 0.018, *p*_sz_ = 5.9 e^-04^) significantly elevated in SZs. Functionally localized time-resolved and dynamic entropy averages are highly correlated with each other; in both cases heightened in blocks involving cognitive control (**Figure 3**), and some blocks of DMN and VIS; relatively lower, among intra-domain blocks, in SC-SC, VIS-VIS and SM-SM. The latter two are blocks that tend to remain consistently strongly cohesive and intercorrelated, making them important to determining cluster membership (**Figure 4**), but evidently not loci of functional reorganization that requires periods of disorder. In the case of *dyn*Ent, DMN-DMN is also among the lower intra-domain blocks (**Figure 3**), indicating that while the DMN is not highly functionally integrated, the disorder is static or gradually manifesting on timescales longer than Δ*t=*1. As in the case of local-global cluster matching (**Figure 4**), the SZ effects on blockwise *tr*Ent and *dyn*Ent (**Figure 3**) directionally accentuate the tendencies in the population averages: SZ has significant negative effects on VIS-VIS time-resolved and dynamic entropy; significant positive effects on VIS-to-other and SC-to-other time-resolved and dynamic entropy. SZ appears to be pushing the general trends toward either order or disorder to greater extremes.

**Figure 3.**
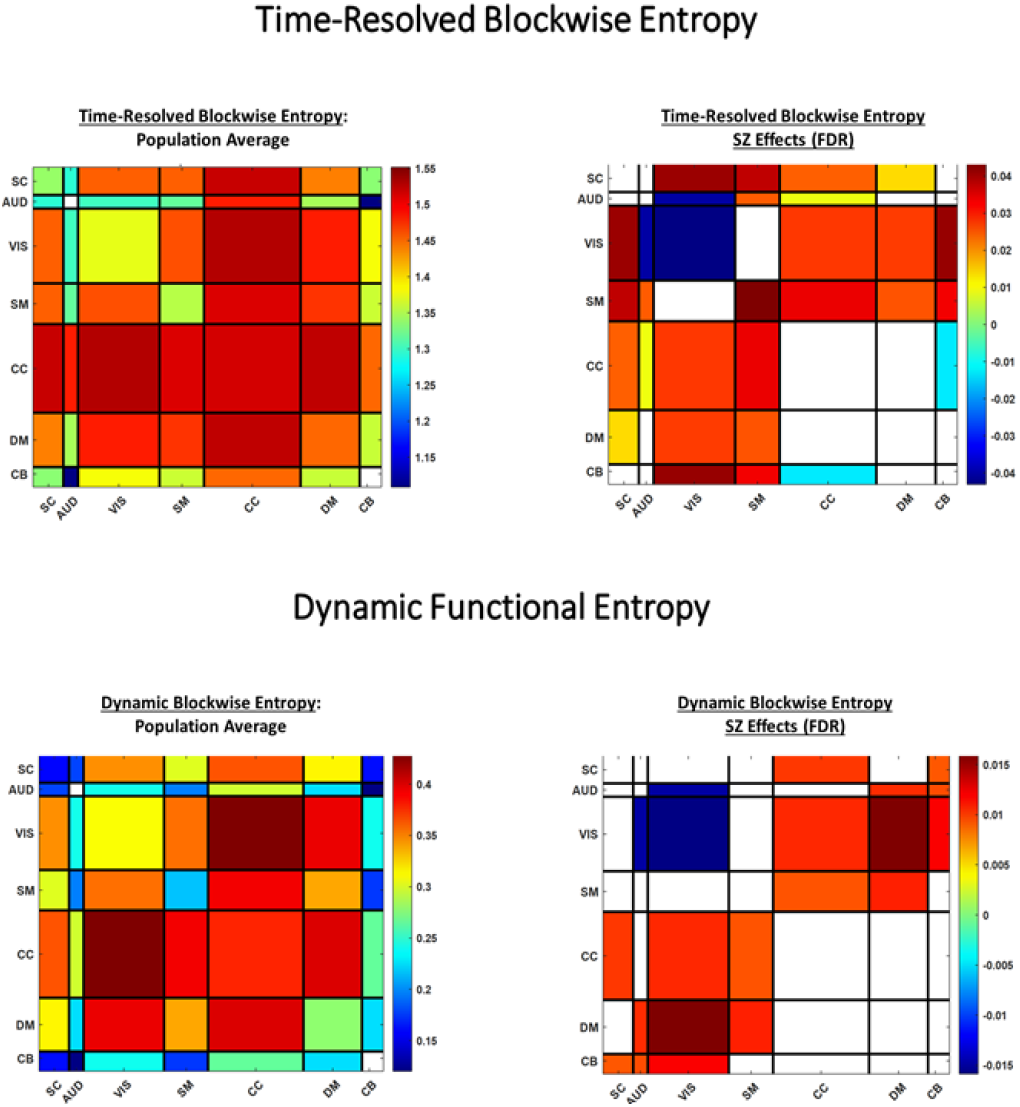
(Top) Time resolved blockwise entopy; (left) CC-SC, CC-VIS, CC-DMN all notably elevated; (right) SZ further suppresses VIS-VIS entropy and elevates VIS-to-other entropy; (Bottom) Dynamic blockwise entropy; population average (left) and SZ effects (right) follow patterns similar to those of *tr*Ent; population averaged DMN-DMN has relatively lower *dyn*Ent than *tr*Ent and SZ effects on DMN-SM *dyn*Ent are relatively stronger.

**Figure 4.**
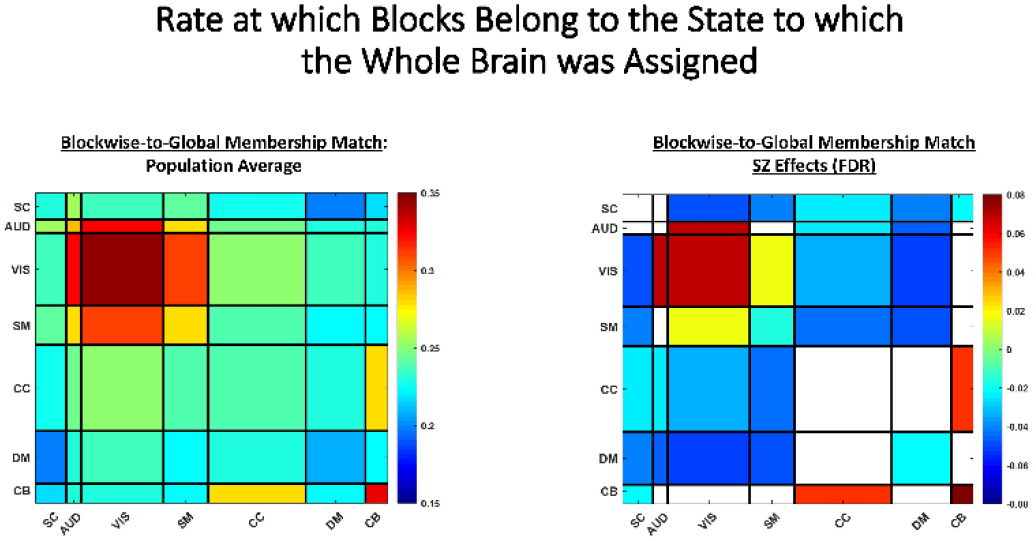
(Left) Domain blocks are assigned membership to the global cluster to which the largest percentage of their individual cells are most proximal. The averaging effect of whole-brain dFNC clustering means that the elementwise match within domain blocks can be close to chance (0.2), with highest local-global match (0.32) in the strongly intercorrelated AUD-VIS-SM (ASVM) block, suggesting thi block is very influential in whole-brain dFNC cluster-based analyses; (Right) non-AVSM blocks are even more poorly matched to global cluster in SZs; AVSM better matched.

Another question is whether time-resolved and/or dynamic entropy are different, on functionally localized terms, immediately preceding a whole-brain transition between global clusters (**Figure 5**). In the case of blockwise *tr*Ent the answer is yes, and the specific domain blocks that gain or lose integrative cohesion depend heavily on the source and destination global state (**Figure 5**). Focusing on Markov loops, i.e., the self-to-self transitions associated with periods dwelling in a fixed global state, we see that the disconnected global state (#5) “tolerates” the highest amount of within-state functional disorder, while staying within either of the highly modularized AVSM-dominant global states (#2 and #4) seems more delicate, requiring less disorder/ more integrative cohesion. Not surprisingly, blockwise *dyn*Ent (**Figure 5**) is notably depressed when remaining in a fixed global state, with the exception of weakly connected global state 5 in which in which occupancy can persist with significantly elevated *dyn*Ent in VIS-to-other domain blocks (**Figure 5**).

**Figure 5.**
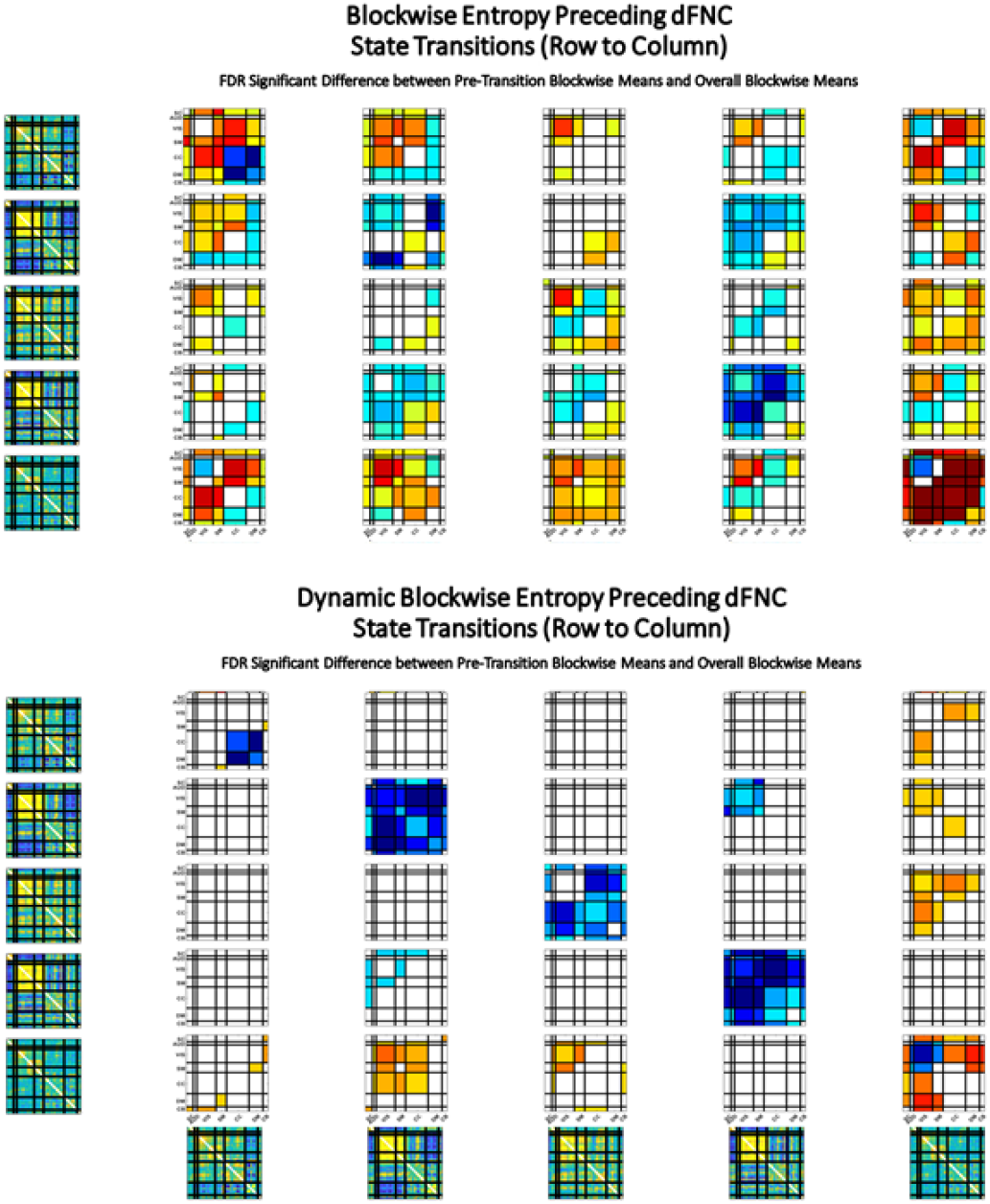
(Top) Significant *t*-statistics from *t*-tests for differences of time-resolved blockwise entropy in timesteps immediately preceding a whole-brain transition from the global state indicated along rows, to the global state indicated below columns (bottom panel) from the overall blockwise mean *tr*Ent; staying in the disconnected global state (row 5) is associated with above average *tr*Ent in many blocks, while remaining in either of the modular, AVSM-dominant global states (rows 2 and 4) is associated with below average *tr*Ent in many blocks; (Bottom) Same display for dynamic blockwise entropy. Notably, with the exception of disconnected state 5, the within-state transitions exhibit pervasively and significantly below average blockwise *dyn*Ent, which is consistent with remaining in a given global state. Dynamic entropy preceding self-transitions within disconnected state 5 is elevated in VIS-to-other blocks, further emphasizing the role of this state in absorbing fluctuating, heterogeneous patterns of functional integration.

## 4. DISCUSSION

We introduced here a framework for integrating functionally localized time-varying connectomic information into a commonly employed pipeline for studying time-varying connectivity at the whole brain level. Initial findings indicate that the degree to which domain blocks within the connectome are reflected in whole-brain cluster centroids is limited: not far from chance for many domain blocks, and most prominent in the highly intercorrelated AVSM block (**Figure 4**). The mismatch outside of the AVSM block is significantly more pronounced in patients than controls. Moreover, using entropy as a metric for (the absence of) integrative functional cohesion, we find that patients exhibit significantly higher whole-brain *tr*Ent, whole-brain *dyn*Ent, blockwise *tr*Ent in the VIS-to-other blocks, and significantly lower blockwise *tr*Ent and *dyn*Ent entropy within the AUD-VIS block (**Figure 3**).

Resolving our findings to timepoints that precede whole-brain state transitions reveals strong patterning in the functionally-localized precursor conditions for global state shifts (**Figure 5**). The case of Markov loops, i.e. self-to-self transitions, provides a tractable space to quickly identify some mechanistically and clinically relevant trends. Mechanistically, we see that *dyn*Ent tends to be lower than average preceding a global self-loop. The exception is global state 5, a weakly connected state in which significant disorder is “tolerated” both functionally and dynamically. In the case of *tr*Ent, we see evidence of relatively cohesive functional integration in the self-loops between modularized AVSM-dominated global states 2 and 4, both more occupied by controls at a whole-brain level. Global state 1, more occupied by patients, is split during self-loops between high *tr*Ent in VIS-to-other blocks, and more functional cohesion in the DMN. Finally, the weakly connected global state 5, more occupied by patients at the whole-brain level, exhibits suppressed functional cohesion in the AUD-VIS block and heightened dissipative disorder in the remainder of the connectome during self-loops.

There is also a clear mapping of characteristic SZ effects on blockwise entropy (**Figure 3**) and the blockwise entropic conditions consistent with remaining in global state 5 (**Figure 5**), a state implicated in the dysconnectivity theory of schizophrenia [3], suggesting that functionally localized dynamic conditions are playing an important role in this theory and its whole-brain evidentiary support.

## 5. ACKNOWLEDGMENTS

This work was supported by NIH grants R01MH123610 and R01MH118695.

